# Expansion, retention and loss in the Acyl-CoA Synthetase *“Bubblegum”* (*Acsbg*) gene family in vertebrate history

**DOI:** 10.1101/288530

**Authors:** Mónica Lopes-Marques, André M. Machado, Raquel Ruivo, Elza Fonseca, Estela Carvalho, L. Filipe C. Castro

**Author notes:** Address correspondence to: Mónica Lopes-Marques & Luís Filipe Costa Castro Interdisciplinary Centre for Marine and Environmental Research (CIIMAR) Av. General Norton de Matos s/n, 4450-208 Matosinhos, Portugal.

## Abstract

Fatty acids (FAs) constitute a considerable fraction of all lipid molecules with a fundamental role in numerous physiological processes. In animals, the majority of complex lipid molecules are derived from the transformation of FAs through several biochemical pathways. Yet, for FAs to enroll in these pathways they require an activation step. FA activation is catalyzed by the rate limiting action of Acyl-CoA synthases. Several Acyl-CoA enzyme families have been previously described and classified according to the chain length of FA they process. Here, we address the evolutionary history of the ACSBG gene family which activates, FA with more than 16 carbons. Currently, two different ACSBG gene families, *ACSBG1* and *ACSBG2*, are recognized in vertebrates. We provide evidence that a wider and unequal *ACSBG* gene repertoire is present in vertebrate lineages. We identify a novel *ACSBG-like* gene lineage which occurs specifically in amphibians, ray finned fish, coelacanths and chondrichthyes named *ACSBG3*. Also, we show that the *ACSBG2* gene lineage duplicated in the Theria ancestor. Our findings, thus offer a far richer understanding on FA activation in vertebrates and provide key insights into the relevance of comparative and functional analysis to perceive physiological differences, namely those related with lipid metabolic pathways.

## 1. Introduction

Lipids represent a complex group of biomolecules present in all living organisms, playing a key role in numerous biological processes, such as inflammatory response, reproduction, biological membranes and energy sourcing and storage. Additionally, they participate in the overall the homoeostasis as signal molecules, cofactors, and endogenous ligands for nuclear receptors (Wall et al., 2010; Robinson and Mazurak, 2013; Grygiel-Górniak, 2014). Aside from sterol lipids, all remaining lipids are obtained from the endogenous elaboration of fatty acid (FA) molecules (Watkins et al., 2007). Yet, for FAs to enroll in any anabolic and catabolic process they require an activation step. Thus, FA activation is a critical rate limiting step of FA metabolism. FA activation was first recognized in 1948 and referred to as “sparking” or “priming” (Grafflin and Green, 1948; Knox et al., 1948). This enzymatic step consists of a two-step thioesterification reaction catalyzed by Acyl-CoA synthetase (ACS), resulting in a thioester with coenzyme A (CoA) (Watkins et al., 2007).

Several ACS involved in FA activation have been previously identified and organized according to the degree of unsaturation and chain length of the FAs favored as substrate: the short-chain ACS-Family (*ACSS*), medium-chain ACS-Family (*ACSM*), long-chain ACS-Family (*ACSL*), very long-chain ACS-Family (*ACSVL*), Bubblegum ACS-Family (*ACSBG*) and ACSFamily (*ACSF*) (Watkins et al., 2007; Soupene and Kuypers, 2008). Although some substrate preference overlap is observed, these enzymes also differ in tissue distribution and subcellular location, an indication of their highly specific role in FA metabolism (Watkins et al., 2007). ACS enzymes have been found to have a wide taxonomic distribution, with homologues ranging from Eubacteria to Plants and Metazoa, a clear indication of their pivotal role in lipid metabolism (Hisanaga et al., 2004; Soupene and Kuypers, 2008).

Despite their wide taxonomic occurrence, the genetic repertoire of ACS has been found to vary, namely in vertebrates (Castro et al., 2012; Lopes-Marques et al., 2013). For example, some studies have disclosed that both ACSL and ACSS gene family composition and function were shaped by events of gene/genome duplication in combination with differential loss. Moreover, multi-genome comparisons across a wide range of vertebrate species revealed novel and previously uncharacterized ACS enzymes (Castro et al., 2012; Lopes-Marques et al., 2013). The present work seeks to build on previous findings and further extend, the knowledge regarding the genetic repertoire and distribution ACS in vertebrates namely the ACS *Bubblegum* (ACSBG) gene family.

ACSBG enzymes, also known as lipidosin, activate FA with C16 to C24 (Moriya-Sato et al., 2000; Steinberg et al., 2000; Pei et al., 2003). Presently, 2 members of the ACSBG gene family have been identified and characterized in mammals, ACSBG1 and ACSBG2 (Pei et al., 2003; Watkins et al., 2007). Similarly, to the previously described ACS enzymes, both ACSBG members display conserved sequence motifs, such as the putative ATP-AMP signature motif for ATP binding (Motif I) and a motif for FA binding, characteristic of the ACS gene family (Motif II) (Moriya-Sato et al., 2000; Watkins et al., 2007). Notably, all known ACS, with the exception of human ACSBG2, contain a highly conserved arginine (Arg-R) in Motif II (Pei et al., 2006). The replacement of this Arg by histidine (His-H) in Human ACSBG2 was found to confer a biphasic pH optimum (pH 6.5 and pH 7.5) to the enzyme, in contrast to the monophasic activity at pH 7-7.5 of the mouse orthologue (Pei et al., 2006). Yet, due to the degree of conservation of both Motifs I and II, these have previously been used to seek and identify potential ACS enzymes (Steinberg et al., 2000; Watkins et al., 2007).

ACSBG enzymes have been suggested to play a significant role in brain development and reproduction (Moriya-Sato et al., 2000; Tang et al., 2001; Pei et al., 2006). Previous reports with the *D. melanogaster* bubblegum mutant (termed bubblegum due to the bubbly appearance of the lamina, a result of neurodegeneration and dilation of the photoreceptor axons) and mouse, associated the disruption of *ACSBG1* to X-linked adrenoleukodytrophy (X-ALD) (Min and Benzer, 1999; Moriya-Sato et al., 2000). X-ALD is characterized by the accumulation of high levels of very long FA in plasma and tissues, accompanied by neurodegeneration (Min and Benzer, 1999; Moriya-Sato et al., 2000; Moser et al., 2002). On the other hand, *ACSBG2* plays an important role in spermatogenesis and testicular development, being associated to male infertility (Zheng et al., 2005; Fraisl et al., 2006). In agreement gene expression of *ACSBG1* is found to be mainly restricted to brain, adrenal gland, gonads, spleen in mouse and human (Moriya-Sato et al., 2000). In contrast, *ACSBG2* showed a more exclusive expression pattern being highly expressed in the testis, followed by medulla and spinal cord (Pei et al., 2006).

Here, using a combination of phylogenetics, comparative genomics and gene expression analysis we deduced the evolutionary history of the *ACSBG* gene family in all major vertebrate lineages. Our findings illustrate the importance of comparative analysis to address the role of adaptive evolution in the shaping of lipid metabolic modifications between lineages.

## 2. Materials & Methods

### 2.1. Database search and phylogenetic analysis

NCBI GenBank release 220 June and release 221 August 2017 and Ensembl release 89 May and release 90 August 2017, databases were searched using tblastn and blastp to recover ACSBG-like sequences using human ACSBG1 (NP_055977) and ACSBG2 (NP_112186) amino acid sequences as query. All major vertebrate lineages such as mammals, birds, reptiles, amphibians, coelacanths, teleost fish, cartilaginous fish and cyclostomes were searched. Additionally, the following invertebrate lineages basal to chordates were also explored cephalochordates, hemichordates, Mollusca (Supplementary material 1).

Our search retrieved 121 ACSBG-like sequences; sharing a minimum 70% pairwise identity with corresponding query sequence for mammals; 60% for bird’s reptiles and amphibians; 50% identity for teleost’s and chondrichthyes and finally 40% identity for invertebrates. The collected sequences were aligned and inspected with partial sequences being removed, leaving a 119 full ORF or near full ORF sequences for phylogenetic analysis (Supplementary material 1). Amino acid sequences were aligned in MAFFT with the L-INS-I method (Katoh et al., 2005; Katoh and Toh, 2008). In the resulting alignment, all columns containing 90% gaps were stripped leaving a total of 787 positions for phylogenetic analysis. A second sequence alignment containing 121 ACSBG-like sequences including the truncated sequences of *Xenopus tropicalis* and *Xenopus leavis* was performed using the same method leaving a total of 788 positions for phylogenetic analysis. Both alignments were then individually submitted to PhyML3.0 server (Guindon et al., 2010), with evolutionary model determined automatically, resulting in the selection JTT+G+I in both cases. The branch support for phylogenetic trees was calculated using aBayes. The resulting trees were visualized and edited in Fig. Tree V1.3.1 available at http://tree.bio.ed.ac.uk/software/figtree/ and rooted with the invertebrate sequences.

### 2.2. Synteny and Paralogy analysis

Using as reference the human and teleost *ACSBG loci*, synteny maps of the genomic neighbourhoods of the *ACSBG1, ACSBG2* and *ACSBG3* gene were assembled in a set of species representative of the major lineages analysed. The following genome assemblies available in NCBI were accessed for *Homo sapiens* - GCF_000001405.33, *Monodelphis domestica* - GCF_000002295.2, *Gallus gallus* - GCF_000002315.4, *Pelodiscus sinensis* - GCF_000230535.1, *X. tropicalis* - GCF_000004195.3, *X. laevis* - GCF_001663975.1, *Latimeria chalumnae -* GCF_000225785.1, *Oryzias latipes* - GCA_000313675.1, *Astyanax mexicanus* - GCA_000372685.2, *Danio rerio* - GCA_000002035.4, *Lepisosteus oculatus* - GCA_000242695.1, *Callorhinchus milii -* GCA_000165045.2, *Branchiostoma floridae* - GCA_000003815.1 and *S. kowalevskii -* GCF_000003605.2. For *Tetraodon nigroviridis* and *Petromyzon marinus* genome assemblies TETRAODON 8.0, Mar 2007 and Pmarinus_7.0, Jan 2011 available in Ensembl were accessed. Paralogy analysis of the *ACSBG locus* was conducted using the reconstructed ancestral chordate genome as described in (Putnam et al., 2008).

### 2.3. RNA isolation and ASCBG tissue expression panel in *X. tropicalis*

*X. tropicalis* (African clawed frog) tissues (brain, skin, heart, liver, spleen, pancreas, kidney, intestine, testis and ovary) were kindly provided by O. Brochain (CNRS, Orsay). Total RNA was purified using the Illustra RNAspin Mini RNA Isolation Kit animal tissues protocol (GE Healthcare) with on-column DNase I digestion. RNA quality was assessed by electrophoresis and its concentration determined using a microplate spectrophotometer (Take 3 and Synergy HT Multi-Mode Microplate Reader, Biotek). First-strand cDNA was synthesized from 250ng RNA using the iScriptTM cDNA Synthesis Kit (Bio-Rad), according to the manufacturer’s instructions.

Forward and reverse primers sets were designed to flank an intron and to avoid genomic DNA amplification. Primers sets were created for the following genes *ACSBG1* - Forward-5′TTTGCCAGGATGTTGGAAGT3′, Reverse-5′AAAGCTTCCACGTGCTCTGT 3′, annealing at 57°; *ACSBG2* - Forward-5′ CTTTTCTGGGGACGTCATGT 3′, Reverse-5′ TTGGAACCTGCTCTTTGAGG 3′, annealing at 55° and *ASCGB3* - Forward-5′ TGCAGTCTTTGCTACGTTGG 3′ reverse-5′ ACAAACAGAGCTCCCCTGTG 3′, annealing at 57°. To assess the quality of *X. tropicalis* cDNA two sets of primers targeting housekeeping genes were included for β-actin – Forward-5’ GGTCGCCCAAGACATCAG3′, Reverse-5′GCATACAGGGACAACACA annealing at 57º and for EEF1A1 – Forward-5′TCGTTAAGGAAGTCAGCACA3′ and Reverse5′CATGGTGCATTTCAACAGAT3′ annealing at 57º. PCR reactions were all performed using 2 μl of *X. tropicalis* cDNA and Phusion® Flash high-fidelity Master Mix (FINNZYMES). PCR parameters were as follows: initial denaturation at 98°C for 10 s, followed by 30 cycles of denaturation at 98°C for 1 s, annealing for 5 s and elongation at 72°C for 10s and a final step of elongation at 72°C for 1 min. PCR products were then loaded onto 2% agarose gel stained with GelRed and run in TBE buffer at 80 V.

### 2.4. ACSBG expression analysis through Rna-Seq

The RNA-Seq analysis was performed using a collection of tissues datasets from seven species Human (*H. sapiens*), mouse (*M. musculus*), chicken (*G. gallus*), western clawed frog (*X. tropicalis*), zebrafish (*D. rerio*), spotted gar (*L. oculatus*) and elephant shark (*C. milii*), available in National Centre for Biotechnology (NCBI) Sequence Read Archive (SRA) (https://www.ncbi.nlm.nih.gov/sra/) (Supplementary material 2). To standardize datasets from different sources, all files were converted to FASTQ file format and sequence quality trimming was performed using Trimmomatic v 0.36 (Bolger et al., 2014). Reads with 36bp in length and an average score of 20 phred were selected for further analysis.

Reference sequences and respective annotation files of each specie were collected from NCBI and Ensembl (Release 89) (Yates et al., 2016) (supplementary material 3). For both elephant shark and western clawed frog the reference sequences for RNAseq mapping were retrieved from NCBI (ftp://ftp.ncbi.nih.gov/genomes/refseq/), while for the remaining species the reference sequences for mapping were retrieved from Ensembl database (ftp://ftp.ensembl.org/pub/release-89/) (Supplementary material 3). Trimmed and groomed reads from each dataset were mapped to their respective reference using Bowtie2 (Langmead and Salzberg, 2012), and the transcript quantification was calculated in transcript per million (TPM), with RSEM v.1.2.31 software (Li and Dewey, 2011). TPM values for each gene were taken as evidence of relative gene expression, low TPM values (< 0.5) were considered unreliable and substituted with zero. To complete this exploratory gene expression analysis, the TPM values were log_2_-transformed after adding a value of one.

## 3. Results and discussion

### 3.1. Database mining and phylogenetic analysis reveals novel members ACSBG gene family

Initial blast searches identified ACSBG-like sequences and recovered a larger than anticipated number of sequence hits. ACSBG1 and ACSBG2-like sequences were found in species from the following vertebrate lineages: mammals, birds, reptiles, amphibians, holostei, coelacantiforms and teleostei. In chondrichthyans and cyclostomes no ACSBG1-like sequences were retrieved. Moreover, an additional uncharacterized ACSBG2-like sequence was also retrieved in some mammalian species. Database searches also recovered a novel set of ACSBG-like sequences in four gnathostome lineages: amphibians (Western clawed frog and African clawed frog) chondrichthyans (elephant shark), holostei (spotted gar), coelacantiforms (coelacanth) and teleostei. However, amphibian sequences were considerably shorter than the remaining ACSBG, thus being excluded from the main phylogenetic analysis.

To disclose the orthology of these various sequences a phylogenetic analysis was performed. The resulting tree topology displays 3 well supported clades in vertebrates. The first group contained all ACSBG1 sequences from mammals, reptiles, birds, amphibians, coelacanths and teleost fish, with no representatives of chondrichtyes and cyclostomes. Besides the ACSBG1 clade, we find a sister clade comprising ACSBG2 sequences. This contains ACSBG2 previously described in mammals (ACSBG2a) and an uncharacterized ACSBG2-like (ACSBG2b) identified in the present work. The tree topology suggests that the both mammalian ACSBG2 sequences are related by a duplication event that took place in the ancestor of Theria. Out grouping the mammalian ACSBG2 sequences we find the ACSBG2 sequences from birds, reptiles, amphibians, coelacanths, chondrichthyes, teleost fish and cyclostomes. Thus, *bona fide* orthologues of ACSBG2 are represented across all major vertebrate classes. Within the third clade, we find a novel uncharacterized group of ACSBG sequences which have sequence representatives in coelacanths, teleost fish, holostei and chondrichthyes. We name this novel sequence ACSBG3. Finally, placed basally to all vertebrate sequences we find invertebrate ACSBG sequences. The present tree topology provides robust indications that the diversification of the ACSBG gene family occurred in the vertebrate ancestor. A second phylogenetic analysis was run separately to include the uncharacterized short *Xenopus sp.* ACSBG sequences (Supplementary material 4). Although the overall tree topology is conserved with the main analysis (Fig. 1), we find that the *Xenopus sp.* ACSBG sequences are placed basally to all vertebrate clades hindering the identification of their orthology. This placing of *Xenopus sp*. ACSBG-like sequences correlates to the highly divergent nature observed in the sequence alignment, with these amphibian sequences being considerably shorter and displaying a poor conservation of the AMP-binding motif (see section 3.3).

**Figure 1:**
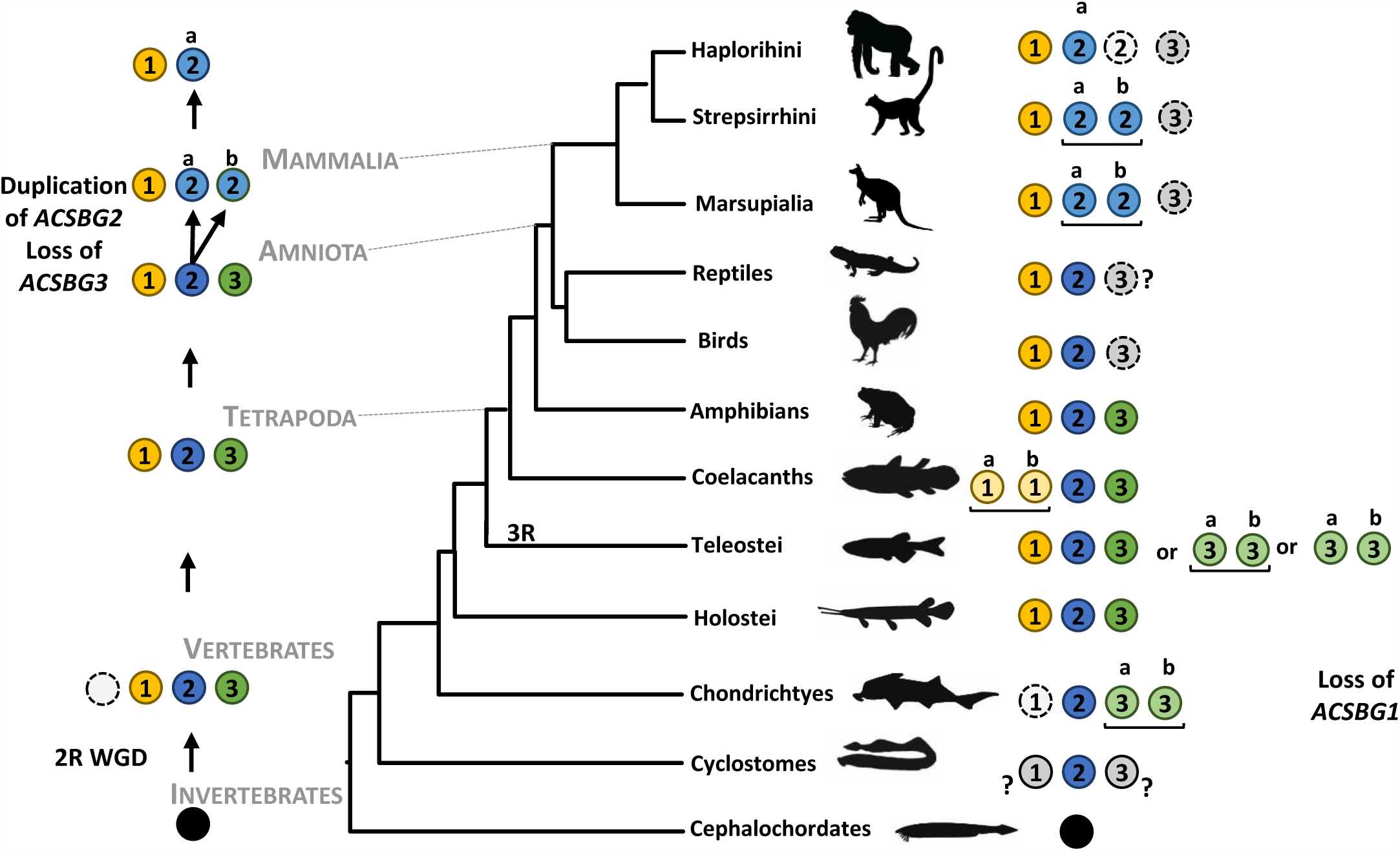
Maximum likelihood phylogenetic analysis of ACSBG amino acid sequences rooted with the invertebrate clade. Numbers at nodes indicate posterior probabilities calculated using aBayes.

Additionally, we find that mammalian ACSBG2a and 2b sequences are placed in long branches in both phylogenetic analysis (Fig.1 and Supplementary material 4) suggesting an accelerated evolution and divergence of these sequences further analysed in section 3.3.

### 3.2. A novel Acsbg gene, Acsbg3, is an ohnolog gone missing in amniotes

Phylogenetic analysis suggests that the ACSBG gene family expanded in the vertebrate ancestor. This time frame coincides with the proposed timing of two round of whole genome duplication (2R WGD) in the ancestral vertebrate approximately 500MYA (Ohno, 1970; Dehal and Boore, 2005). Yet, while it is generally accepted that all gnathostomes underwent the 2R-WGD, the extent of these genome duplications in cyclostomes still remains a matter of debate (Smith and Keinath, 2015).

To complement the phylogenetic analysis, validate events of gene duplication/loss, resolve the orthology of the *Xenopus sp.* uncharacterized ACSBG and the origin of ACSBG3 sequences, the genomic *locus* of each *Acsbg* gene was examined in a set of representative species (Fig. 2). Comparative synteny analysis of the ACSBG1 *locus* reveals a high degree of conservation of neighbouring gene families throughout all the analysed lineages (Fig.2A). *ACSBG1* is localized in human chromosome 15, being flanked by gene families such as the *IDH3A, CIB2, WDR61* and *CRABP1*. These flanking gene families are also present in the vicinity of ACSBG1 *locus* in the all analysed lineages (Fig. 2A). In the case of *C. milii* although no *ACSBG1* gene was found, synteny analysis reveals that the *locus* organization is conserved, suggesting gene loss in this lineage (Fig. 2A). Regarding the cyclostomes, extensive blast searches did not retrieve an ACSBG1 sequence; synteny analysis uncovered a fragmented *locus* segregated into at least two distinct scaffolds. Therefore, the absence of this gene in cyclostomes may be attributed to poor genome coverage or to gene loss (Fig. 2A).

**Figure 2:**
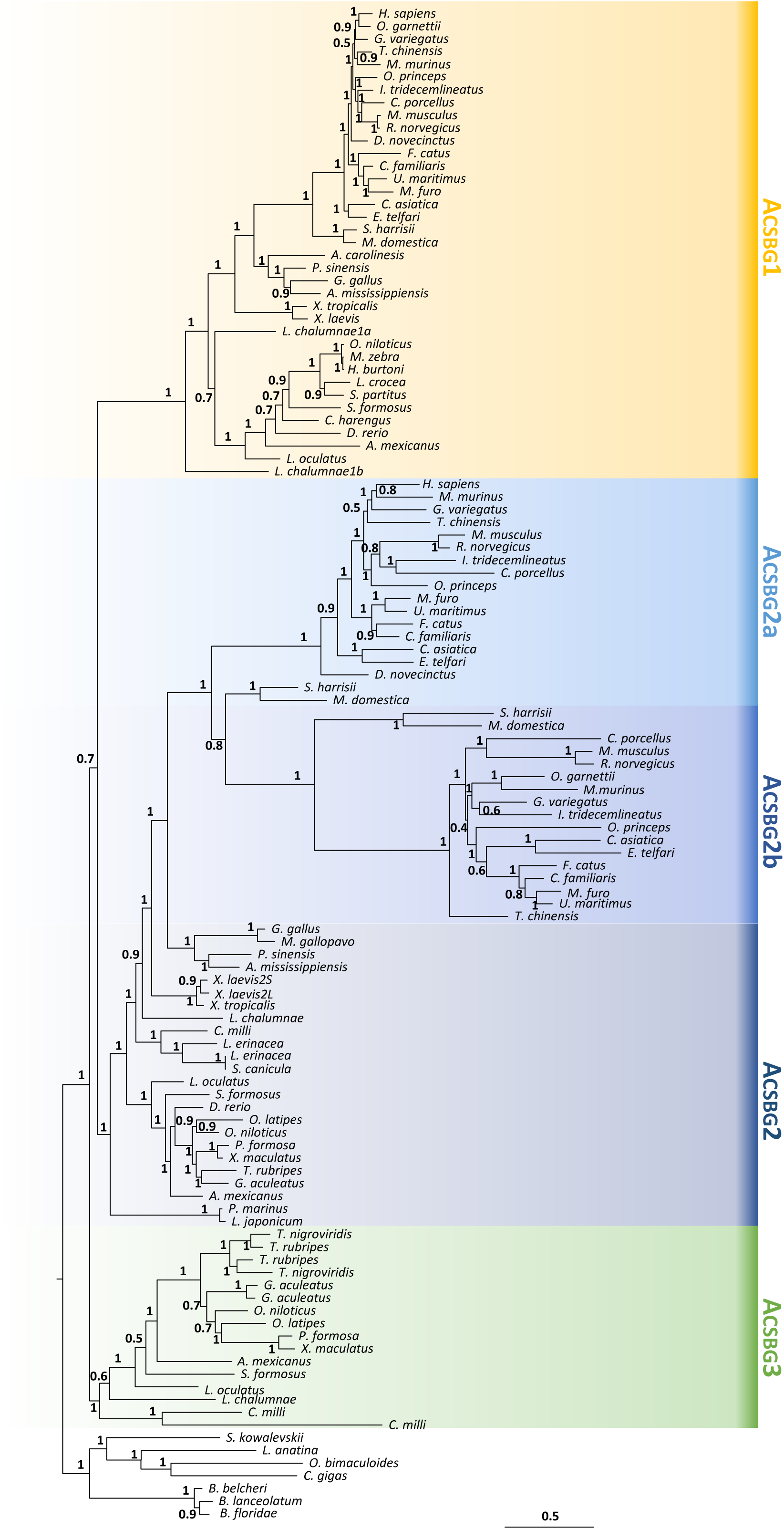
Comparative genomic maps of vertebrate *ACSBG1* (A) *ACSBG2* (B) and *ACSBG3* (C and D) gene *loci.* Paralogy analysis and invertebrate genomic maps of ACSBG (E and F).

The human ACSBG2 gene resides in chromosome 19 and is flanked by the following gene families: *RFX2, RANBP3, MLLT1* and *ACER1.* The *locus* architecture is conserved in all species analysed. In cyclostomes, the *ACSBG2 locus* is disjointed and distributed among several scaffolds thus the absence of ACSBG2 in cyclostomes remains similarly to ACSBG1 unresolved (Fig. 2B). In the case of mammalian duplicates, we find that *ACSBG2a* and *ACSBG2b* are located side by side in the *M. domestica.* Synteny analysis of this *locus* in other mammals presenting both *ACSBG2a* and *2b* (data not shown) is coincident with the observation for *M. domestica* supporting the hypothesis that *ACSBG2a* and *2b* arose through tandem duplication in the ancestor of therian mammals, with ACSBG2b being later lost in Haplorhini. Finally, we find that in the *ACSBG3 locus*, despite the lesser conservation, some neighbouring gene families such as *HINT2, SPAG8, RGP1* and *GBA2* are preserved in the majority of the analysed lineages. Using these conserved neighbouring gene families, the corresponding *locus* was mapped in birds and mammals to address the loss of *ACSBG3* in these lineages (Fig. 2C). ACSBG3 is also absent in reptiles and the analysis of the *locus* revealed that it is fragmented in various species examined (*Anolis carolinesis*, *Thamnophis sirtalis*, *Alligator mississippiensis* and *Chrysemys picta*), hindering the validation of ACSBG3 loss in this lineage.

We next investigated the synteny maps for the single ACSBG *locus* from two invertebrate cephalochordates. Here we find that the *B. floridae locus* retains a conserved gene family arrangement, namely with the presence of *HERC1-like* gene, whose orthologue is found in the vicinity of vertebrate *ACSBG1* (Fig. 2 A and D indicated in red). Similarly, the hemichordate *S. kowalevskii* also displays a conserved neighbouring gene family, *CHRNA3* with the vertebrate orthologue placed in the *ACSBG1 locus* (Fig. 2A and D indicated in red). Finally, to address the hypothesis that ACSBG gene expansion took place with the 2R WGD the location of ACSBG and neighbouring genes (with described paralogues underlined genes in Fig.2A, B, C and D) were mapped to the predicted ancestral paralogons as described by Putnam *et al* 2008 (Putnam et al., 2008). Next, ancestral paralogons were mapped back to the same ancestral linkage group, LG2 (Putnam et al., 2008) indicating that the *ACSBG loci* are related by genome duplication, strongly suggesting that vertebrate ACBG diversity arose with the 2R WGD (Fig. 2 F).

### 3.3. Sequence analysis and gene expression

To further characterize the novel *ACSBG2b* and *ACSBG3* a sequence alignment was performed to inspect the typical ACS enzyme motifs (Watkins et al., 2007). The analysis of this alignment revealed that the predicted AMP-binding domain (Motif I Fig. 3A and 3B), a highly conserved motif in all ACS enzymes from bacteria to humans (Black et al., 1997; Steinberg et al., 2000; Weimar et al., 2002; Karan et al., 2003), is conserved in the vast majority of the sequences collected with the exception of the novel ACSBG2b (Fig.3B) and the ACSBG3 sequence in *Xenopus sp.*(see Supplementary material 5 for alignment of the full 121 sequences). An indication that residues within Motif I play a fundamental role in ACS catalytic activity was found in previous studies were the mutation of residues within Motif I (positions 1, 2, 4, 5 and 10) in *E. coli* considerably reduced the catalytic activity, while the replacement of residues 1 and 5 in *S. cerevisiae* resulted in a minor reduction of enzymatic activity (Fig. 3A grey arrows) (Weimar et al., 2002; Zou et al., 2002). Thus, the low conservation of this motif in the mammalian-specific ACSBG2b strongly suggests that these enzymes may show an alternative function or *modus operandi*. In the analysis of the *Xenopus sp.* ACSBG3, we find that this motif differs from the remaining ACSBG3 identified, being disrupted with the deletion of 3bp (Supplementary material 5). Again, this observation suggests an alternative role for the enzyme given that AMP binding is essential for FA activation. Regarding Motif II, also known as the ACS signature-motif and proposed to be involved in acyl chain length specificity (Black et al., 1997), we find that again ACSBG2b displays a divergent sequence when compared to the remaining ACSBG analysed here. Interestingly, we observe that ACSBG2b presents an Asparagine residue (Asn-N) instead of the highly conserved Arginine (Fig. 3A and B black arrow). Notably, human ACSBG2a harbours a Histidine in this position, representing the single case described to date of an ACSBG without an Arginine (Pei et al., 2006). Reverse mutation of the Histidine within Motif II in human ACSBG2a showed that this residue assumes a critical role in determining the optimal pH for this enzyme (Pei et al., 2006). Additionally, this replacement (Asn) is only observed for placental mammals, with marsupials retaining the conserved Arginine (Fig 3 B). Next, Motif V (KXX(R,K) is a conserved motif found in several members of the ACS enzymes families and contains a conserved K-Lys demonstrated to be essential for the catalytic function of ACS in *S. enterica* propionyl-coA synthetase and ACS activity of murine ACSF2 (Horswill and Escalante-Semerena, 2002). Here we find that Motif V is conserved in all recovered ACSBG sequences with the exception of *Xenopus sp* ACSBG3 due to the short nature of these sequences (see Supplementary material 5). Finally, the Motifs III and IV, identified by Hisanaga *et al* 2008 (Hisanaga et al., 2004), are found to be conserved in the majority of analysed sequences, with the exception of a conservative replacement in Motif IV in ACSBG2b. The highly conserved Histidine is replaced by biochemically similar residue, tyrosine, having a minor or no predicted impact.

**Figure 3:**
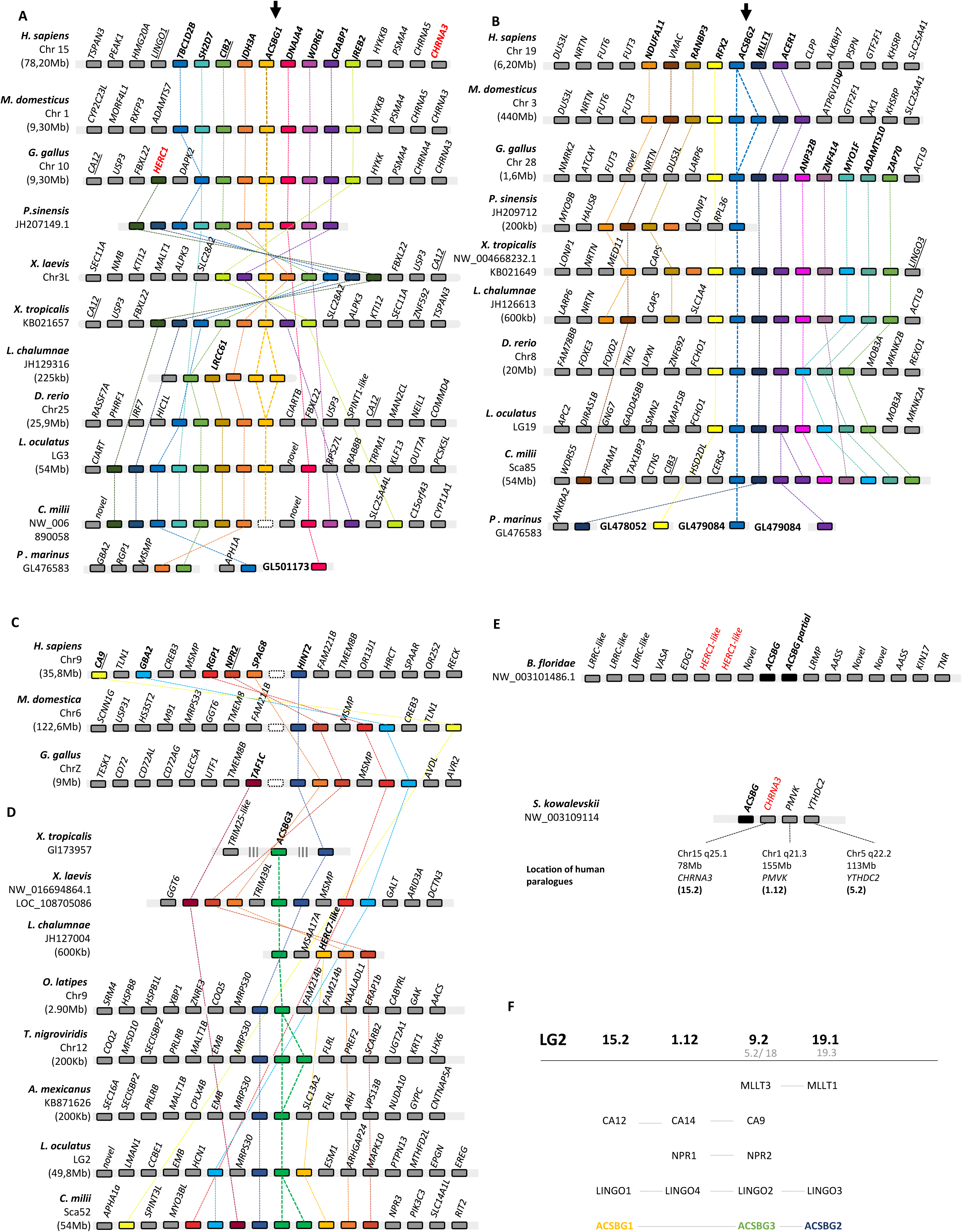
Sequence alignment and ACS Motif analysis. **A**-Sequence logo graphs of the consensus sequences of all ACSBG sequences recovered excluding mammal specific ACSBG2b acyl-coenzyme A sequences, totalizing 104 sequences. **B**-Sequence logo graphs of the consensus sequences of all mammalian specific ACSBG2b (17 sequences). Overall height of the stack reflects the degree of conservation, the height of each letter represents relative frequency of a given residue in a specific position. Black arrow highlights the highly conserved arginine residue (Pei et al., 2006) and corresponding position in ACSBG2b **C**-Heatmap of the relative expression of *ACSBG1 ACSBG2* and *ACSBG3* obtained from RNA-seq analysis and visualized using Matrix2png (Pavlidis and Noble, 2003). **D-** Tissue expression profile of *X. tropicalis* ACSBG genes.

In an attempt to infer the function of the newly identified *ACSBG2b* and *ACSBG3*, and address the retention of these genes in a restricted number of lineages, we next performed an expression analysis using available RNA-Seq SRAs (Fig. 3C). Similarly, to previous reports (Moriya-Sato et al., 2000; Tang et al., 2001; Pei et al., 2006), relative expression profiles reveal that *ACSBG1* expression is mainly limited to brain and gonads, with the exception of *D. rerio* for which the liver stands as the main expression site. On the other hand, the expression profile of non-mammalian vertebrate *ACSBG2* was found to be more extensive than in mammals (Pei et al., 2006) with expression detected in all analysed tissues of *C. milii*, *D. rerio*, *L. oculatus*, *G. gallus*, and *X. tropicalis*. Interestingly, the expression analysis of mammalian specific duplicates *ACSBG2a* and *ACBG2b* shows a confined expression of the duplicates essentially in testis, with a relatively low expression of *ACSBG2a* detected in human kidney. Regarding the gene expression profile of *ACSBG3 in X. tropicalis, L. oculatus*, and in *ACSBG3a* of *C. milli*, we find a localized and high relative expression in ovary and in testis. Semi-quantitative PCR expression analysis of ACSBG3 from *X. tropicalis* is in accordance with *in silico* RNAseq analysis (Fig. 3D), with ACSBG1 expression confined to brain and testis, while ACSBG2 is detected all tissues except ovary and finally ACSBG3 being restricted to testis and brain (Fig. 3D).

High expression of *ACSBG3* in gonads is indicative that this enzyme may play an important role in reproduction similarly to the role of *ACSBG2* (Pei et al., 2006). Finally, for *ACSBG3* in *C. milii* no expression was detected in any of the analysed tissues.

### 3.4. Evolutionary history of ACSBG gene family

Using a multi-comparative approach, including database searches, phylogenetic and synteny analysis we have uncovered a larger than anticipated genetic repertoire of *ACSBG* genes in vertebrates. We find that the initial expansion of the *ACSBG* gene family from which arose *ACSBG1 ACSBG2* and *ACSBG3* is coincident with the 2R WGD, with representative gene orthologues present in several gnathostome lineages (Fig. 4). The detailed analysis of the *ACSBG* gene repertoire revealed a differential retention of *ACSBG3*, with this paralogue being lost in birds, mammals and possibly in reptiles, while being retained in teleosts, amphibians and chondrichthyes. The identification of additional ACSBG enzymes in teleosts correlates with previous studies, where differential paralogue retention led to the maintenance of extra ACS enzyme paralogues, namely *ACSL2* and *ACSS1b* in teleosts (Fraisl et al., 2006; Wall et al., 2010). The preservation of duplicated genes is often observed when the corresponding transcript, in this case ACS, is in high demand (Zhang, 2003). Thus, one may hypothesize that the preservation of additional ACS duplicates in teleosts is a means to fulfil a high demand of FA activation given that FA oxidation is considered to be the main energy source in this lineage (Tocher, 2003). Finally, further duplications were observed in the ancestor of mammals with the tandem duplication of *ACSBG2* and also in specific lineages such as the *ACSBG3* in *C. milii* and *T. nigroviridis* and *ACSBG1 L. chalumnae* (Fig. 4).

**Figure 4:**
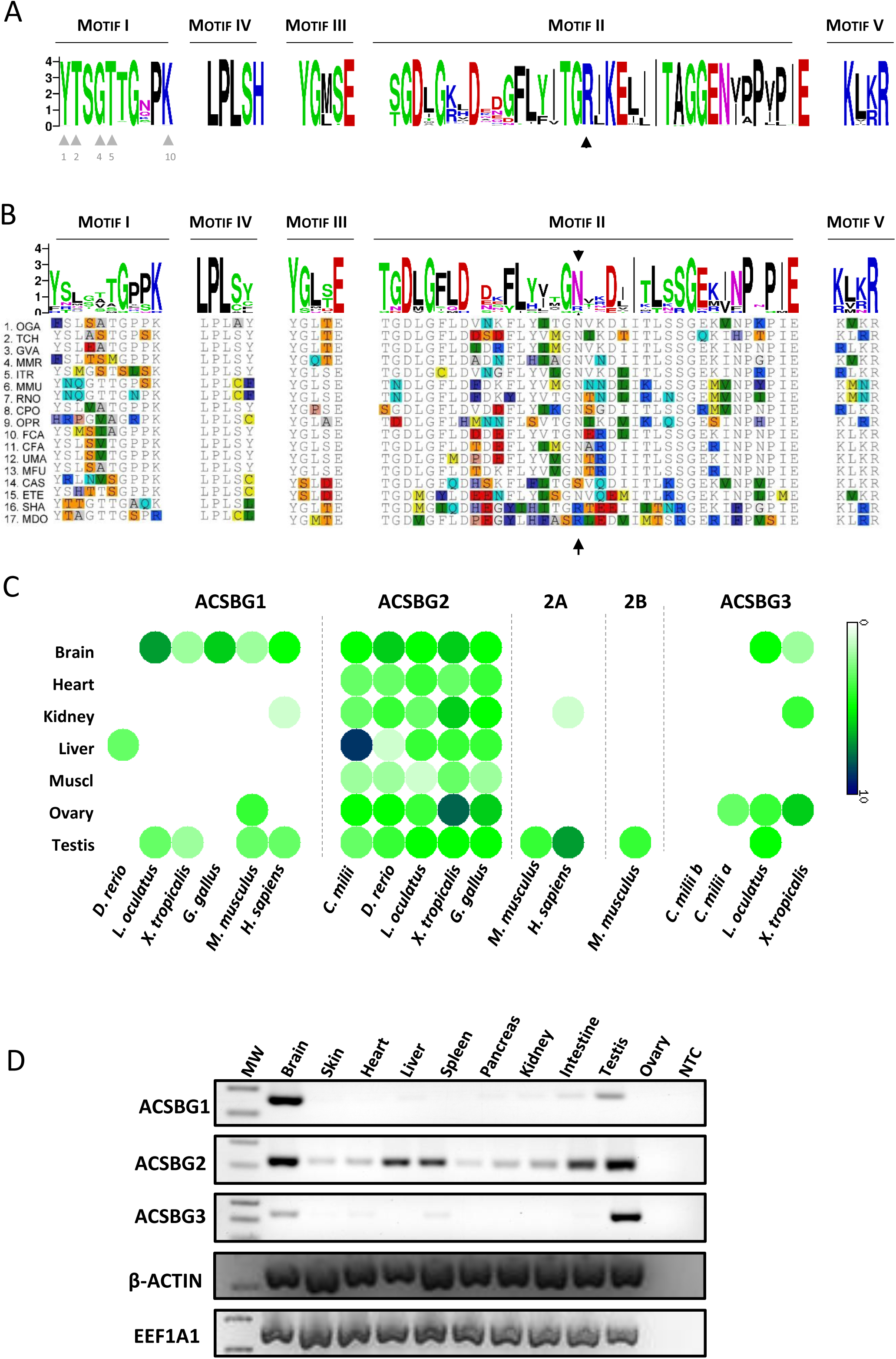
Proposed evolutionary history of the *ACSBG* gene family in vertebrates. Yellow corresponds to *ACSBG1* blue *ACSBG2* and green *ACSBG3*. Grey full lined circles with question marks indicate unknown or unresolved if gene is present, grey circles dashed lined indicate gene loss. Black line under genes indicates tandem duplication.

## 4. Conclusion

Our findings suggest that FA activation metabolic modules, including the ACSBG gene family, have significantly diversified upon vertebrate radiation as a consequence of genome duplication, lineage specific duplication and losses.

## Acknowledgments

This work was supported by Project INNOVMAR—Innovation and Sustainability in the Management and Exploitation of Marine Resources (reference NORTE-01-0145-FEDER-000035, within Research Line NOVELMAR/INSEAFOOD/ECOSERVICES), supported by North Portugal Regional Operational Programme (NORTE 2020), under the PORTUGAL 2020 Partnership Agreement, through the European Regional Development Fund (ERDF). We acknowledge Fundação para a Ciência e a Tecnologia for the support to EF (SFRH/BD/79305/2011).

**Supplementary material 1:** Table containing ACSBG sequences accession numbers.

**Supplementary material 2:** Accession numbers of the RNAseq files retrieved for expression analysis.

**Supplementary material 3:** Genome and GTF files retrieved from Ensemble database (Release 89) and Transcriptome files retrieved from NCBI used on this study and accession numbers of reference genes.

**Supplementary material 4:** Complementary phylogenetic analysis of ACSBG sequences including amphibian uncharacterized truncated ACSBG-like sequences indicated in red.

**Supplementary material 5:** Motif sequence alignment of the full dataset used 121 ACSBG sequences. Red box highlights mammal specific ACSBG2b.

