## Supplementary Materials for "Expansion, retention and loss in the Acyl-CoA Synthetase *“Bubblegum”* (*Acsbg*) gene family in vertebrate history"

**Supplementary material 2.** Accession numbers of the RNAseq files retrieved for expression analysis.

| Human | Library | SRA Run | Read length |
|---|---|---|---|
| BRAIN | PE | ERR315432 | 100 |
| HEART | PE | ERR315328 | 100 |
| KIDNEY | PE | ERR315468 | 100 |
| LIVER | PE | ERR315451 | 100 |
| MUSCLE | PE | SRR1617452 | 100 |
| OVARY | PE | ERR315402 | 100 |
| TESTIS | PE | ERR315456 | 100 |
| **Mouse** | **Library** | **SRA Run** | **Read length** |
| BRAIN | PE | SRR5171101 | 100 |
| HEART | PE | SRR5171076 | 100 |
| KIDNEY | PE | SRR5171094 | 100 |
| LIVER | PE | SRR5171078 | 100 |
| MUSCLE | PE | SRR1556527 | 100 |
| OVARY | PE | SRR5171100 | 100 |
| TESTIS | PE | SRR5171084 | 100 |
| **Chicken** | **Library** | **SRA Run** | **Read length** |
| BRAIN | PE | SRR924542 | 100 |
| HEART | PE | SRR924549 | 100 |
| KIDNEY | PE | SRR924553 | 100 |
| LIVER | PE | SRR924555 | 100 |
| MUSCLE | PE | SRR924559 | 100 |
| OVARY | PE | SRR924548 | 100 |
| TESTIS | PE | SRR924547 | 100 |
| **Western clawed frog** | **Library** | **SRA Run** | **Read length** |
| BRAIN | PE | SRR579560 | 75 |
| HEART | PE | SRR579563 | 75 |
| KIDNEY | PE | SRR579562 | 75 |
| LIVER | PE | SRR579561 | 75 |
| MUSCLE | PE | SRR579564 | 75 |
| OVARY | SE | SRR943352 | 100 |
| TESTIS | SE | SRR943353 | 100 |
| **Spotted gar** | **Library** | **SRA Run** | **Read length** |
| BRAIN | PE | SRR1524250 | 100 |
| HEART | PE | SRR1524252 | 100 |
| KIDNEY | PE | SRR1524255 | 100 |
| LIVER | PE | SRR1524254 | 100 |
| MUSCLE | PE | SRR1524253 | 100 |
| OVARY | PE | SRR1524259 | 100 |
| TESTIS | PE | SRR1524260 | 100 |
| **Zebrafish** | **Library** | **SRA Run** | **Read length** |
| BRAIN | PE | SRR1524238 | 100 |
| HEART | PE | SRR1524240 | 100 |
| KIDNEY | PE | SRR1524243 | 100 |
| LIVER | PE | SRR1524242 | 100 |
| MUSCLE | PE | SRR1524241 | 100 |
| OVARY | PE | SRR1524248 | 100 |
| TESTIS | PE | SRR1524249 | 100 |
| **Elephant Shark** | **Library** | **SRA Run** | **Read length** |
| BRAIN | PE | SRR514109 | 75 |
| HEART | PE | SRR514107 | 75 |
| KIDNEY | PE | SRR514105 | 75 |
| LIVER | PE | SRR513760 | 75 |
| MUSCLE | PE | SRR514104 | 75 |
| OVARY | PE | SRR513759 | 75 |
| TESTIS | PE | SRR513757 | 75 |
