## Supplementary Materials for "Expansion, retention and loss in the Acyl-CoA Synthetase *“Bubblegum”* (*Acsbg*) gene family in vertebrate history"

**Supplementary material 3**

**Table A:** Genome and GTF files retrieved from Ensemble database (Release 89) and Transcriptome files retrieved from NCBI used on this study.

| Organisms | Genome / Transcriptome Files | GTF FILE | Download date | Source |
|---|---|---|---|---|
| **Mouse** | Mus_musculus.GRCm38.dna.toplevel.fa | Mus_musculus.GRCm38.89.gtf | 29/07/2017 | Ensemble |
| **Spotted gar** | Lepisosteus_oculatus.LepOcu1.dna.toplevel.fa | Lepisosteus_oculatus.LepOcu1.89.gtf | 29/07/2017 | Ensemble |
| **Zebrafish** | Danio_rerio.GRCz10.dna.toplevel.fa | Danio_rerio.GRCz10.89.gtf | 29/07/2017 | Ensemble |
| **Human** | Homo_sapiens.GRCh38.dna.toplevel.fa | Homo_sapiens.GRCh38.89.gtf | 29/07/2017 | Ensemble |
| **Chicken** | Gallus_gallus.Gallus_gallus-5.0.dna.toplevel.fa | Gallus_gallus.Gallus_gallus-5.0.89.gtf | 29/07/2017 | Ensemble |
| **Western clawed frog** | GCF_000004195.3_Xenopus_tropicalis_v9.1_rna.fna | Xenopus_tropicalis_Gene_Transcript_Map.txt* | 29/07/2017 | Ncbi |
| **Elephant Shark** | GCF_000165045.1_Callorhinchus_milii-6.1.3_rna.fna | Callorhinchus_milii_Gene_Transcript_Map.txt* | 29/07/2017 | Ncbi |

*Built in lab from *rna.fna file and batch entrez of NCBI (https://www.ncbi.nlm.nih.gov/sites/batchentrez )

**Table B:** Accession numbers of reference genes used map and quantify RNAseq reads.

| Organisms | ACSBG1 | ACSBG2 | ACSBG2a | ACSBG2b | ACSBG3 |
|---|---|---|---|---|---|
| **Mouse** | ENSMUSG00000032281 | - | ENSG00000130377 | ENSMUSG00000024209 | - |
| **Spotted gar** | ENSLOCG00000014820 | ENSLOCG00000000968 | - | - | ENSLOCG00000011989 |
| **Zebrafish** | ENSDARG00000062077 | ENSDARG00000004094 | - | - | - |
| **Human** | ENSG00000103740 | - | ENSG00000130377 | - | - |
| **Chicken** | ENSGALG00000003286 | ENSGALG00000001749 | - | - | - |
| **Western clawed frog** | Gene Id:100158596 | Gene Id:100158533 | - | - | Gene Id:100494473 |
| **Elephant Shark** | - | Gene Id: 103183485 | - | - | Gene Id:103181008<br>Gene Id:103181022 |
