## Supplementary Materials for "Expansion, retention and loss in the Acyl-CoA Synthetase *“Bubblegum”* (*Acsbg*) gene family in vertebrate history"

**Supplementary material 4**: Complementary phylogenetic analysis of ACSBG sequences including amphibian uncharacterized truncated ACSBG-like sequences indicated in red.

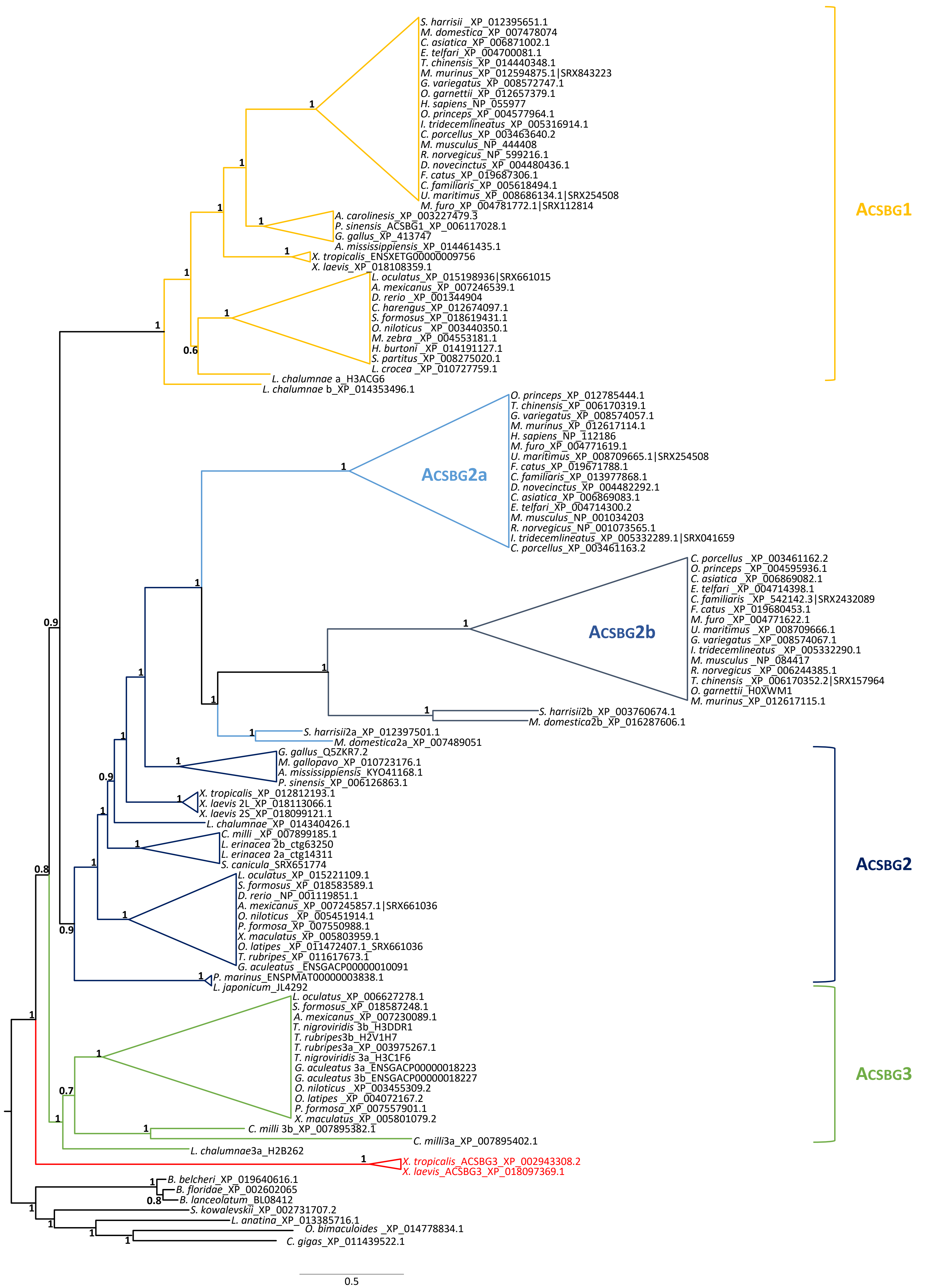
