## Supplementary Materials for "Expansion, retention and loss in the Acyl-CoA Synthetase *“Bubblegum”* (*Acsbg*) gene family in vertebrate history"

**Supplementary material 5:** Motif sequence alignment of the full dataset used 121 ACSBG sequences. Red box highlights mammal specific ACSBG2b.

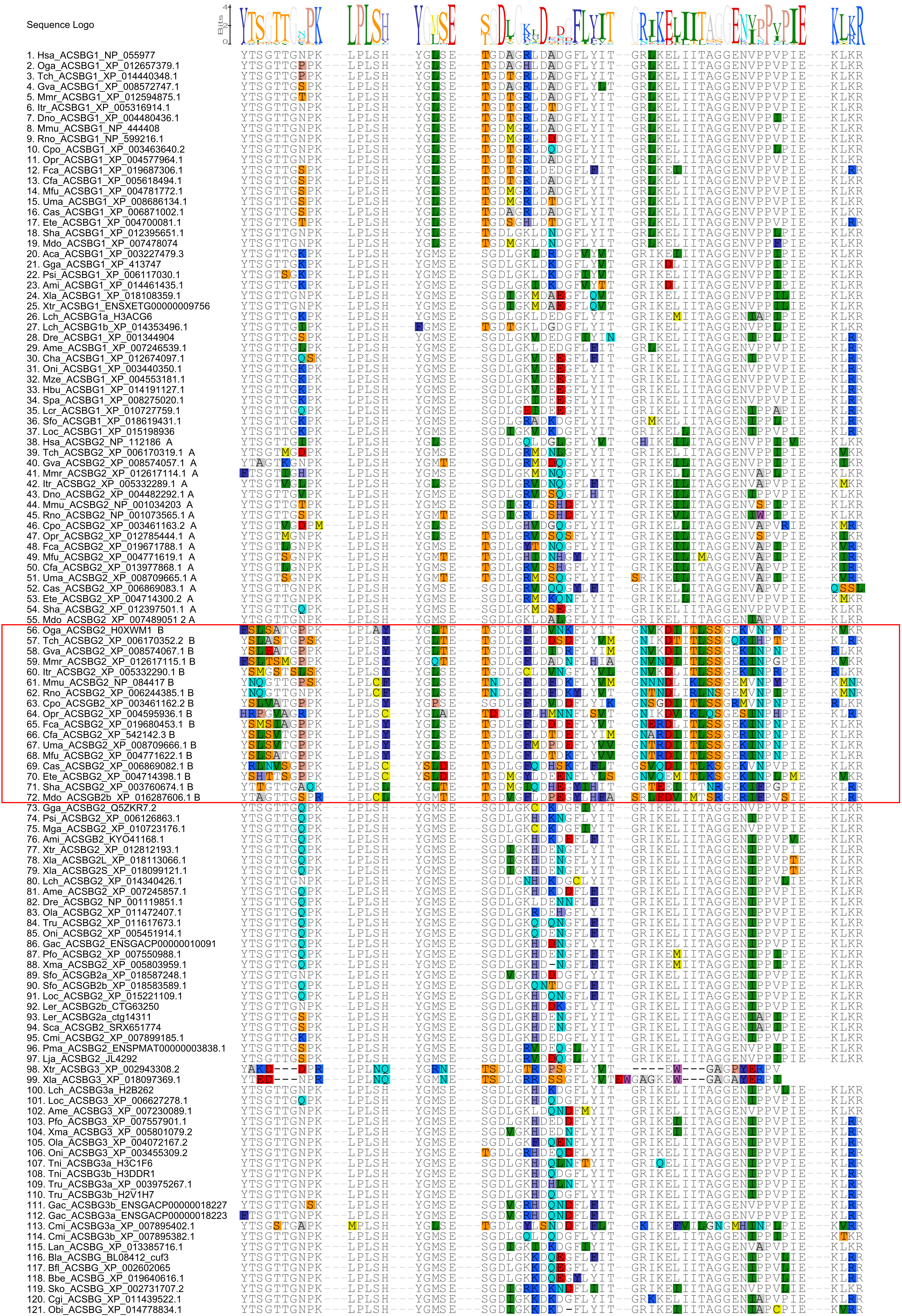
