## Supplementary Materials for "Expansion, retention and loss in the Acyl-CoA Synthetase *“Bubblegum”* (*Acsbg*) gene family in vertebrate history"

**Supplementary material 1**: ACSBG sequences accession numbers.

| Specie | ACSBG1 | ACSBG2 | | ACSBG3 |
|---|---|---|---|---|
| ***Homo sapiens*** | NP_055977 | NP_112186 | Lost | |
| ***Otolemur garnettii*** | XP_012657379.1 | - | H0XWM1 | |
| ***Ictidomys tridecemlineatus*** | XP_005316914.1 | XP_005332289.1 | XP_005332290.1 | |
| ***Microcebus murinus*** | XP_012594875.1 | XP_012617114.1 | XP_012617115.1 | |
| ***Tupaia chineses*** | XP_014440348.1 | XP_006170319.1 | XP_006170352.2 | |
| ***Cavia porcellus*** | XP_003463640.2 | XP_003461163.2 | XP_003461162.2 | |
| ***Mus musculus*** | NP_444408 | NP_001034203 | NP_084417 | |
| ***Rattus norvegicus*** | NP_599216.1 | NP_001073565.1 | XP_006244385.1 | |
| ***Felis catus*** | XP_019687306.1 | XP_019671788.1 | XP_019680453.1 | |
| ***Canis lupus familiaris*** | XP_005618494.1 | XP_013977868.1 | XP_542142.3 | |
| ***Ursus maritimus*** | XP_008686134.1 | XP_008709665.1 | XP_008709666.1 | |
| ***Mustela putorius furo*** | XP_004781772.1 | XP_004771619.1 | XP_004771622.1 | |
| ***Ochotona princeps*** | XP_004577964.1 | XP_012785444.1 | XP_004595936.1 | |
| ***Galeopterus variegatus*** | XP_008572747.1 | XP_008574057.1 | XP_008574067.1 | |
| ***Dasypus novemcinctus*** | XP_004480436.1 | XP_004482292.1 | | |
| ***Echinops telfairi*** | XP_004700081.1 | XP_004714300.2 | XP_004714398.1 | |
| ***Chrysochloris asiatica*** | XP_006871002.1 | XP_006869083.1 | XP_006869082.1 | |
| ***Monodelphis domestica*** | XP_007478074 | XP_007489051 | XP_016287606.1 | |
| ***Sarcophilus harrisii*** | XP_012395651.1 | XP_012397501.1 | XP_003760674.1 | |
| ***Anolis carolinensis*** | XP_003227479.3 | | | |
| ***Pelodiscus sinensis*** | XP_006117030.1 | XP_006126863.1 | | |
| ***Alligator mississippiensis*** | XP_014461435.1 | KYO41168.1 | | |
| ***Gallus gallus*** | XP_413747 | Q5ZKR7.2 | | |
| ***Meleagris gallopavo*** | | XP_010723176.1 | | |
| ***Xenopus laevis*** | XP_018108359.1 | XP_018113066.1<br>XP_018099121.1 | | XP_018097369.1 |
| ***Xenopus tropicalis*** | ENSXETG00000009756 | XP_012812193.1 | | XP_002943308.2 |
| ***Latimeria chalumnae*** | H3ACG6 | XP_014340426.1 | | H2B262 |
| ***Astyanax mexicanus*** | XP_007246539.1 | XP_007245857.1 | | XP_007230089.1 |
| ***Clupea harengus*** | XP_012674097.1 | | | |
| ***Takifugu rubripes*** | | XP_011617673.1 | | XP_003975267.1<br>H2V1H7 |
| ***Gasterosteus aculeatus*** | | ENSGACP00000010091 | | ENSGACP00000018223<br>ENSGACP00000018227 |
| ***Xiphophorus maculatus*** | | XP_005803959.1 | | XP_005801079.2 |
| ***Poecilia formosa*** | | XP_007550988.1 | | XP_007557901.1 |
| ***Tetraodon nigroviridis*** | | | | H3C1F6<br>H3DDR1 |
| ***Oryzias latipes*** | | XP_011472407.1 | | XP_004072167.2 |
| ***Danio rerio*** | XP_001344904 | NP_001119851.1 | | |
| ***Oreochromis niloticus*** | XP_003440350.1 | XP_005451914.1 | | XP_003455309.2 |
| ***Maylandia zebra*** | XP_004553181.1 | | | |
| ***Haplochromis burtoni*** | XP_014191127.1 | | | |
| ***Larimichthys crocea*** | XP_010727759.1 | | | |
| ***Stegastes partitus*** | XP_008275020.1 | | | |
| ***Scleropages formosus*** | XP_018619431.1 | XP_018583589.1 | | XP_018587248.1 |
| ***Lepisosteus oculatus*** | XP_015198936 | XP_015221109.1 | | XP_006627278.1 |
| ***Callorhinchus milii*** | | XP_007899185.1 | | XP_007895382.1<br>XP_007895402.1 |
| ***Leucoraja erinácea*** | | CTG63250<br>CTG14311 | | |
| ***Scyliorhinus canicula*** | | SRX651774 | | |
| ***Petromyzon marinus*** | | ENSPMAT00000003838.1 | | |
| ***Lethenteron japonicum*** | | JL4292 | | |
| | **INVERTEBRATES** | | | |
| ***Saccoglossus kowalevskii*** | XP_002731707.2 | | | |
| ***Branchiostoma floridae*** | XP_002602065 | | | |
| ***Branchiostoma lanceolatum*** | BL08412 | | | |
| ***Branchiostoma belcheri*** | XP_019640616.1 | | | |
| ***Lingula anatina*** | XP_013385716.1 | | | |
| ***Crassostrea gigas*** | XP_011439522.1 | | | |
| ***Octopus bimaculoides*** | XP_014778834.1 | | | |
